# Escape from nonsense mediated decay associates with anti-tumor immunogenicity

**DOI:** 10.1101/823716

**Authors:** Kevin Litchfield, James Reading, Emilia Lim, Hang Xu, Po Liu, Maise AL-Bakir, Sophia Wong, Andrew Rowan, Sam Funt, Taha Merghoub, Martin Lauss, Inge Marie Svane, Göran Jönsson, Javier Herrero, James Larkin, Sergio A. Quezada, Matthew D. Hellmann, Samra Turajlic, Charles Swanton

**Affiliations:** Cancer Evolution and Genome Instability Laboratory, The Francis Crick Institute, 1 Midland Rd, London NW1 1AT, UK; Cancer Immunology Unit, Research department of Haematology, University College London Cancer Institute, Paul O’Gorman Building, 72 Huntley Street, London, WC1E 6BT, UK; Thoracic Oncology Service, Division of Solid Tumor Oncology, Department of Medicine, Memorial Sloan Kettering Cancer Center, Weill Cornell Medical College, and Parker Center for Cancer Immunotherapy, 885 2nd Avenue, New York, NY 10017, USA; Division of Oncology and Pathology, Department of Clinical Sciences Lund, Faculty of Medicine, Lund University, Scheelegatan 2, Medicon Village, 22185 Lund, Sweden; Center for Cancer Immune Therapy, Department of Hematology, Herlev Ringvej 75, 2730, Herlev, Denmark, Department of Oncology, Copenhagen University Hospital Herlev, Herlev Ringvej 75, 2730, Herlev, Denmark; University College London Cancer Institute, Bill Lyons Informatics Centre, London, WC1E 6DD, UK; Renal and Skin Units, The Royal Marsden Hospital, London, SW3 6JJ, UK; Cancer Research UK Lung Cancer Centre of Excellence, University College London Cancer Institute, Paul O’Gorman Building, 72 Huntley Street, London, WC1E 6BT, UK; Department of Medical Oncology, University College London Hospitals, 235 Euston Rd, Fitzrovia, London, United Kingdom, NW1 2BU, UK

**Keywords:** nonsense mediated decay, neoantigen, mutation, insertions, deletions, indels, immunogenicity, check-point inhibitors, immunotherapy, melanoma

## Abstract

Frameshift insertion/deletions (fs-indels) are an infrequent but potentially highly immunogenic mutation subtype. Although fs-indel transcripts are susceptible to degradation through the non-sense mediated decay (NMD) pathway, we hypothesise that some fs-indels escape degradation and lead to an increased abundance of tumor specific neoantigens, that are highly distinct from self. We analysed matched DNA and RNA sequencing data from TCGA, and five separate melanoma cohorts treated with immunotherapy. Using allele-specific expression analysis we show that expressed fs-indels were enriched in genomic positions predicted to escape NMD, and associated with higher protein expression, consistent with degradation escape (“NMD-escape”). Across four independent cohorts, fs-indel NMD-escape mutations were found to be significantly associated with clinical benefit to checkpoint inhibitor (CPI) therapy (P_meta_=0.0039), a stronger association than either nsSNV (P_meta_=0.073) or fs-indel (P_meta_=0.064) count. NMD-escape mutations were additionally shown to have independent predictive power in the “low-TMB” setting, and may serve as a biomarker to rescue patients judged ineligible for CPI based on overall TMB, but still with a high chance of response (low-TMB cohort: NMD-escape-positive % clinical benefit=53%, NMD-escape-negative % clinical benefit=16%, P=0.0098). Furthermore, in an adoptive cell therapy (ACT) treated cohort, NMD-escape mutation count was the most significant biomarker associated with clinical benefit (P=0.021). Analysis of functional T-cell reactivity screens from recent personalized vaccine and CPI studies shows direct evidence of fs-indel derived neoantigens eliciting patient anti-tumor immune response (n=15). We additionally observe a subset of fs-indel mutations, with highly elongated neo open reading frames, which are found to be significantly enriched for immunogenic reactivity in these patient studies (P=0.0032). Finally, consistent with the potency of NMD-escape derived neo-antigens and ongoing immune-editing, NMD-escape fs-indels appear to be under negative selective pressure in untreated TCGA cases. Given the strongly immunogenic potential, and relatively rare nature of NMD-escape fs-indels, these alterations may be attractive candidates in immunotherapy biomarker optimisation and neoantigen ACT or vaccine strategies.

## Introduction

Tumor mutation burden (TMB) is associated with response to immunotherapy across multiple tumor types, and therapeutic modalities, including checkpoint inhibitors (CPIs) and cellular based therapy (1–8) (9, 10). Although TMB is a clinically relevant biomarker, there are clear opportunities to refine the molecular features associated with response to immunotherapy. In particular, the primary hypothesis about TMB as an immunotherapy biomarker relates to the fact that somatic variants are able to generate tumor specific neoantigens. However, the vast majority of mutations appear to have no immunogenic effect. For example, although hundreds of high affinity neoantigens are predicted in a typical tumor sample, peptide screens routinely detect T cell reactivity against only a few neoantigens per tumor (11). Additionally, the oligoclonal T cell expansions commonly reported in responders to CPI favour the hypothesis that a restricted number of neoantigens mediate anti-tumor immune responses (6). Finally, immunopeptidome profiling via mass spectrometry has similarly identified only a few neoantigens effectively presented on human leucocyte antigen per tumor sample (12). Detailed analysis of TMB to identify the true underlying subsets of mutations driving immunogenicity may substantially optimise biomarker accuracy and improve therapeutic targeting of neoantigens.

We have previously shown that frame shift insertion/deletions (fs-indels) are infrequent (pan-cancer median = 4 per tumor) but a highly immunogenic subset of somatic variants (13). Fs-indels can produce an increased abundance of tumor specific neoantigens with greater mutant-binding specificity. We found that fs-indels are associated with improved response to checkpoint inhibitor (CPI) therapy (13) and may be attractive candidates for therapeutic personalised tumor vaccines. However, fs-indels cause premature termination codons (PTCs) and are susceptible to degradation at the messenger RNA level through the process of non-sense mediated decay (NMD). NMD normally functions as a surveillance pathway to protect eukaryotic cells from the toxic accumulation of truncated proteins. We hypothesized that a subset of fs-indels may escape NMD degradation, and which when translated contribute substantially to directing anti-tumour immunity.

The NMD process is only partially efficient. The canonical NMD model dictates that a mutation triggering a PTC, will escape degradation if located downstream of the last exon junction complex. Therefore, NMD efficiency is intimately linked to sequence position, with reduced efficiency found in the: i) last gene exon, ii) penultimate exon within 50 nucleotides of the 3’ exon junction, and iii) first exon within the first 200 nucleotides of coding sequence (CDS). These rules only partially explain the variance in NMD efficiency however, and an estimated 27% remains unexplained across all genes, increasing to 71% for dosage compensated genes (i.e. genes where copy number deletion is compensated with upregulation of the remaining allele) (14).

Based on these rules, ^~^30% of fs-indels across cancers are predicted to escape NMD (15). Fs-indel mutations escaping NMD have been shown to be an abundant source of expressed neoantigen protein in microsatellite instable (MSI) tumors and to correlate with high levels of CD8 infiltration (16). In addition, targeted inhibition of NMD has been shown to strongly supress tumor growth (17). Taken together these data suggest fs-indels escaping NMD are rare but may be disproportionally immunogenic. Indeed, recent work in parallel with our own from Lindeboom et al. (18) has elegantly demonstrated that NMD escaping fs-indels strongly associate with improved response to CPI therapy. To test this hypothesis further and provide independent validation, we quantified NMD efficiency via allele specific fs-indel detection in paired DNA and RNA sequencing data. We applied this pipeline to four independent cohorts of melanomas treated with CPI, one melanoma adoptive cell therapy cohort, and conducted further NMD analysis in personalized tumor vaccine studies. For further comparison, we also examined non-immunotherapy treated cases from the cancer genome atlas (TCGA).

## Results

### Detection of NMD-escape mutations

Expressed frameshift indels (fs-indels) were detected using paired DNA and RNA sequencing, with data processed through an allele specific bioinformatics pipeline (**Fig. 1A**). Across all processed TCGA samples (n=453, see methods for cohort details) a median of 4 fs-indels were detected per tumor (range 0-470), of which mutant allele expression was detected in a median of 1 per tumor (range 0-94). Thus, expressed fs-indel mutations were present at relatively low frequency and abundance. In fact, 49.6% of samples profiled had zero expressed fs-indel mutations detected. Exon positions were annotated for expressed fs-indels (n=1,840), and compared to non-expressed fs-indels (*i.e*. mutant allele present in DNA, but not in RNA) (n=8,691). Expressed fs-indels were enriched for mutations in last exon positions (odds ratio versus non expressed fs-indels = 1.92, 95% confidence interval [1.69-2.19], P<2.2×10^−16^), as well in penultimate exons within 50 nucleotides of the 3’ exon junction (3’EJ) (OR=1.92 [1.69-2.19], P=4.4×10^−9^). By contrast, expressed fs-indels were depleted in middle exon locations (OR=0.54 [0.48-0.60], P<2.2×10^−16^), and no significant change was detected either way for penultimate exon, >50 nucleotides from the 3’EJ, mutations (**Fig. 1B**). These exon positions are highly consistent with known patterns of NMD-escape, as previously established (14). No depletion/enrichment was detected for first exon (within first 200bp of CDS) position mutations, likely due to the small absolute numbers. Next we considered RNA variant allele frequency (VAF) estimates for expressed fs-indels, and found them to be highest for last (median=0.32), penultimate <=50bp of the 3’EJ (0.32), penultimate >50bp of the 3’EJ (0.27) and first (0.26) exon positions, with middle exon alterations having the lowest value (0.19) (**Fig. 1C**, P<2.2×10^−16^). Finally, we obtained protein expression data from the cancer proteome atlas (19), for 223 proteins across 453 tumors, which overlapped with the DNA/RNAseq processed cohort. Intersecting samples with both an fs-indel gene mutation(s), and matched protein expression data, we compared the protein levels of expressed (n=40) versus non expressed fs-indels (n=96). Protein abundance was found to be significantly higher for expressed fs-indels (P=0.018, **Fig. 1D**). Taken collectively, these results suggest that expressed fs-indels are (at least partially) escaping NMD and being translated to the protein level. Expressed fs-indels are here after referred to as NMD-escape, and non-expressed fs-indels as NMD-competent.

**Figure 1.**
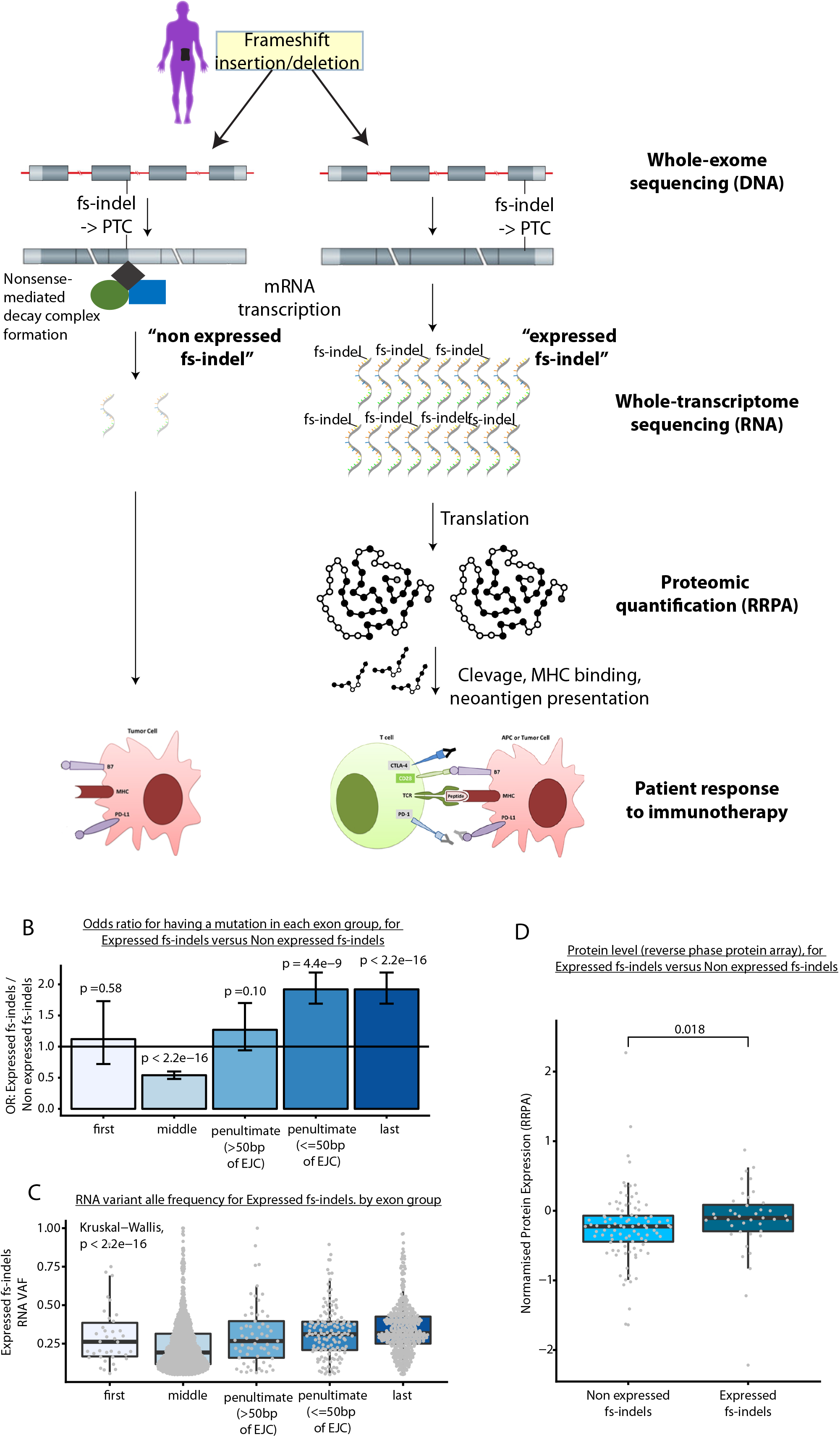
Panel A shows an overview of study deisgn and methodological approach. The left hand side of the panel shows a fs-indel triggered premature termination codon, which falls in a middle exon of the gene, a position associated with efficient non-sense mediated decay (NMD). The right hand side of the panel shows a fs-indel triggered premature termination codon, which falls in the last exon of the gene, a position associated with bypassing NMD. Panel B shows the odds ratio (OR), between expressed fs-indels and non-expressed fs-indels, for falling into either first, middle, penultimate or last exon positions. Odds ratios and associated p-values were calculated using Fisher’s Exact Test. Coloring is used arbitrarily to distinguish groups. Error bars denote 95% confidence intervals of OR estimates. Panel C shows variant allele frequencies for expressed fs-indels by exon group position. Kruskal-Wallis test was used to test for a difference in distribution between groups. Panel D shows protein expression levels for non-expressed, versus expressed, fs-indel mutations. Two-sided Mann Whitney U test was used to assess for a difference between groups.

### NMD-escape mutation burden associates with clinical benefit to immune checkpoint inhibition

To assess the impact of NMD-escape mutations on anti-tumor immune response, we assessed the association between NMD-escape mutation count and CPI clinical benefit in four independent melanoma cohorts with matched DNA and RNA sequencing data: Van Allen et al. (n=33, anti-CTLA-4 treated), Snyder et al. (n=21, anti-CTLA-4 treated), Hugo et al. (n=25, anti-PD-1 treated) and Riaz et al. (n=24, anti-PD-1). For each sample, mutation burden was quantified based on the following classifications: i) TMB: all non-synonymous SNVs (nsSNVs), ii) expressed nsSNVs, iii) fs-indels, and iv) NMD-escape expressed fs-indels. Each mutation class was tested for an association with clinical benefit (**Fig. 2a**). In the pooled meta-analysis of the four melanoma cohorts with both WES and RNAseq (total n=103), nsSNV, expressed nsSNV and fs-indel counts were higher in patients experiencing clinical benefit, but with non-significant p-value (meta-analysis across all cohorts, P_meta_=0.073, P_meta_=0.19 and P_meta_=0.064 respectively) (**Fig. 2a**). NMD-escape mutation count however showed a statistically significant association with clinical benefit (P_meta_=0.0039) (**Fig. 2a**). For clarity, we note sample sizes utilised here are smaller than previously reported, since only a subset of cases had both matched DNA and RNA sequencing data available, and that nsSNV and fs-indel measures are significant in the full datasets. Patients with one or more NMD-escape mutation had higher rates of clinical benefit to immune checkpoint blockade compared to patients with no NMD-escape mutations: 56% versus 12% (Van Allen et al.), 57% versus 14% (Snyder et al.), 75% versus 35% (Hugo et al.) and 64% versus 30% (Riaz et al.) (**Fig. 2b**). We additionally assessed for evidence of correlation between TMB and NMD-escape metrics, and found only a weak correlation between the two variables (r=0.23). And in multivariate logistic regression analysis, we tested both variables together in a joint model to assess for independent significance (n=103, study ID was also included as a model term to control for cohort specific factors), and NMD-escape mutation count was found to independently associate with CPI clinical benefit (P=0.0087), whereas TMB did not reach independent significance (P=0.20). Finally to investigate a potential association in other tumor types, NMD-escape analysis was conducted in a CPI treated metastatic urothelial cancer cohort (n=23 cases) (20). Previous analysis in this study found that neither TMB, predicted neoantigen load nor expressed neoantigen load, were associated with CPI clinical benefit (20). Similarly, here we found no evidence of an association between NMD-escape count and clinical benefit (P=1.0), possibly due to small sample size, lower mutational load lower in this cohort (TMB=^~^0-5 missense SNVs/megabase, as compared to ^~^10.0 in a larger recently published cohort (9)), or lower response rates in general in metastatic urothelial cancer. For completeness, the NMD-escape CPI meta-analysis was repeated to include the above bladder data, together with the four melanoma cohorts, and the association remains significant (P_meta_=0.012).

**Figure 2.**
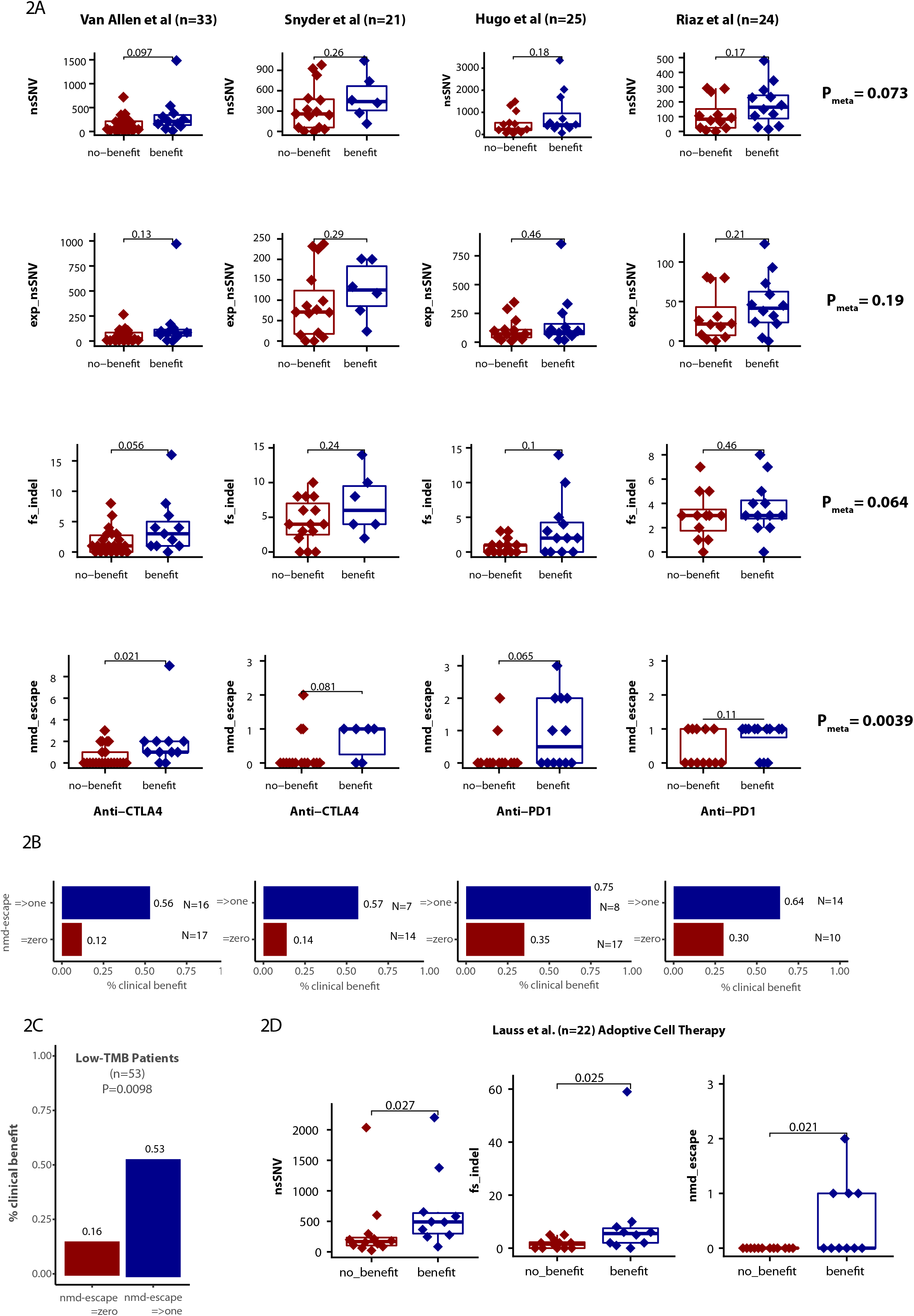
Panel A shows four melanoma checkpoint inhibitor (CPI) treated cohorts, split into groups based on “no-clinical benefit” (dark red) or “clinical benefit” (dark blue) to therapy. Four metrics are displayed per cohort: (top row) TMB non-synonymous SNV count, (second row) expressed non-synonymous SNV count, (third row) frameshift indel count and (fourth row) NMD-escape mutation count. In the first column is the Van Allen et al. anti-CTLA4 cohort, second column is the Snyder et al. anti-CTLA4 cohort, the third column is the Hugo et al. anti-PD1 cohort and the fourth column is Riaz er al. anti-PD1 cohort. Far right are meta-analysis p-values, for each metric across the four cohorts, showing the association with clinical benefit from CPI treatment. Two-sided Mann Whitney U test was used to assess for a difference between groups. Meta-analysis of results across cohorts was conducted using the Fisher method of combining *P* values from independent tests. Panel B shows the % of patient with clinical benefit from CPI therapy, for patients with => 1 NMD-escape mutation (dark blue) and zero NMD-escape mutations (light blue). Panel C shows the combined set of CPI treated patients (across all 4 studies) split to make a low-TMB cohort (nsSNV count < 217, the median value across all cohorts, approximately equivalent to 10 mutations/Mb). The % clinical benefit rates are shown for patients with zero and =>one NMD-escape mutation. Panel D shows non-synonymous SNV count, frameshift indel count and NMD-escape mutation count, compared in an adoptive cell therapy treated cohort.

### NMD-escape mutation burden offers predictive power in low-TMB (nsSNVs) patients

In a clinical scenario where TMB is implemented to stratify patients for CPI therapy, patients with low TMB tumors may be not recommended for CPI treatment. It is known however that some low-TMB tumors can respond to CPI therapy, and we reasoned that NMD-escape mutation count may offer independent predictive power in the low-TMB setting to “rescue” patients who may have a higher chance of response. To investigate this we split the population of CPI treated patients to make a low-TMB cohort (nsSNV count < 217, the median value across all cohorts, approximately equivalent to 10 mutations/Mb), which comprised n=53 patients (all four studies combined). In this cohort NMD-escape mutation count was significantly associated with clinical benefit to CPI (P=0.013), whereas nsSNV count was not (P=0.19). Patients with one or more NMD-escape mutation retained a relatively high rate of clinical benefit from CPI at 53%, compared to 16% for patients with zero NMD-escape events (Odds Ratio = 5.8, 95% confidence interval [1.4 – 27.9], P=0.0098, (**Fig. 2c**). This suggests a potential utility for NMD-escape mutation measurement in tumors with low overall TMB scores.

### NMD-escape mutation burden associates with clinical benefit to adoptive cell therapy

To further investigate the importance of NMD-escape mutations in directing anti-tumor immune response, we analysed matched DNA and RNA sequencing data from patients with melanoma (n=22) treated with adoptive cell therapy (ACT) (10). TMB ns-SNVs (P=0.027), fs-indels (P=0.025) and NMD-escape count (P=0.021) were all associated with clinical benefit from therapy (**Fig. 2d**). All patients with NMD-escape count ≥ 1 experienced clinical benefit (n=4, 100%), compared to 33% (6/18) of patients who had no NMD-escape mutations, further highlighting the potential strong immunogenic effect from just a single NMD-escape mutation.

### Direct evidence of NMD-escape neoantigens

While of translational relevance and clinical utility, biomarker associations do not directly isolate specific neoantigens driving anti-tumor immune response. Accordingly, we obtained data from two anti-tumor personalised vaccine studies, and one CPI study, in which immune reactivity against specific neopeptides had been established by functional assay of patient T cells (21–23). Across these three studies, 15 different fs-indel mutations generated peptides that were functionally validated as eliciting immune reactivity (**Table S1**); thus at a proof of concept level the ability of fs-indels to elicit anti-tumor immune response has been established. Across these same studies, four fs-indel derived neoantigens had also undergone functional screening, but were found to be non-immunogenic (**Table S1**). Although limited by a small sample size, we note that immunogenic fs-indel mutations (n=15) had a significantly longer neoORF length (median=27 amino acids) than screened, but non-immunogenic, fs-indel mutations (n=4, median=5 amino acids, P=0.0032) (**Fig. 3A**). We additionally note several fs-indel mutations with highly elongated neoORF length (termed super neoORF (SNORF) mutations) were detected (neoORF=>50 amino acids, n=5), and these were restricted to the immunogenic group. The number of peptides screened is likely to be a confounding factor in these comparisons (i.e. a longer neoORF allows more unique peptides to be utilised for immunization), however this also highlights the inherent advantage of SNORF events. In the context of SNORF mutations, we next considered redundancy in HLA allele binding, based on the hypothesis that SNORF events (and indeed fs-indels in general) would generate peptides capable of binding to a broader spectrum of patient HLA-alleles. This is likely to be of particular importance in the context loss of heterozygosity at the HLA locus (LOHHLA), a mechanism used by tumor cells to achieve immune evasion. For example considering class I alleles, LOHHLA is known to occur such that one or two HLA-alleles become lost (24), however loss of all six HLA alleles would be unfavourable to the cancer cell, due to global loss of antigen presentation and resulting attraction of natural killer (NK) cell activity (25). We analysed previously published TCGA neoantigen prediction data (26), and note that 56% of nsSNV mutations generate peptides that bind to just one HLA-allele, with only 44% generating peptides than bind to multiple alleles (**Fig. 3B**). By contrast, 17% of fs-indel mutations generate peptides which bind against a single HLA-allele, and instead the majority (83%) bind to multiple HLA-alleles (**Fig. 3B**). In fact ^~^10% of fs-indel mutations (likely SNORF events) were found to generate peptides binding against all six HLA class I alleles, compared to 0.3% of nsSNV mutations (**Fig. 3B**).

**Figure 3.**
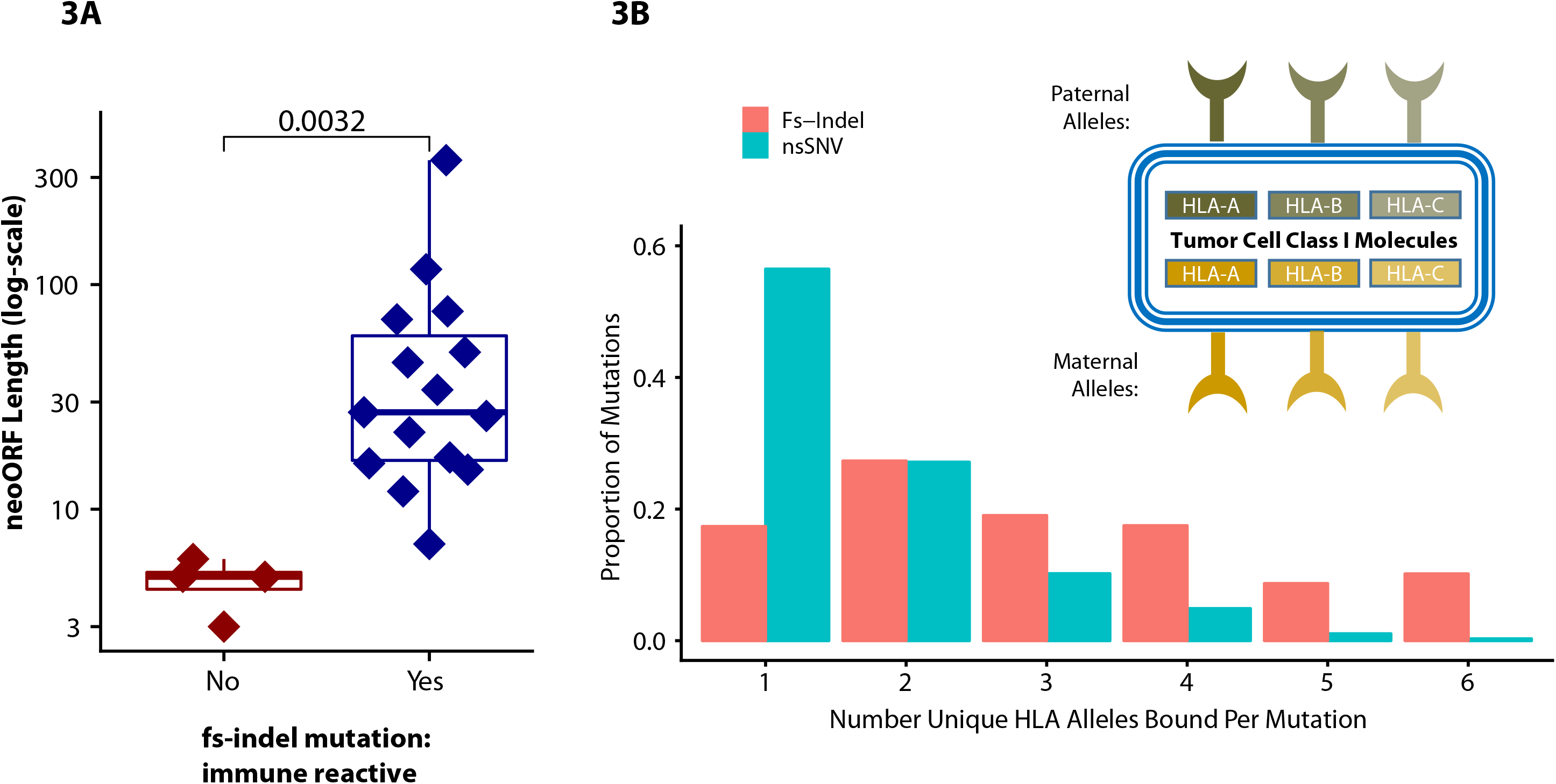
Panel A shows the neo open reading frame (neoORF) length, from functional T-cell reactivity screening data from recent personalized vaccine and CPI studies. In dark red are lengths from fs-indel peptides which were non-reactive, in dark blue are lengths from fs-indel peptides which were T cell reactive. Two-sided Mann Whitney U test was used to assess for a difference between groups. Panel B shows the number of unique class I HLA alleles that individual nsSNV and fs-indel mutations were found to bind against in pan-cancer TCGA data.

### NMD-escape mutations show evidence of negative selection and associate with improved overall survival

Next, we assessed for evidence of selective pressure against NMD-escape mutations, which may reflect the potential to generate native anti-tumor immunogenicity. In additional to potential immunogenic selective pressure, fs-indels have also previously been reported to be under functional selection (15) due to their loss of protein function effect. To account for this, we used stop-gain SNV mutations as a benchmark comparator, as these variants have equivalent functional impact but no immunogenic potential (*i.e*. loss of function but no neoantigens generated). Furthermore, the rules of NMD apply equally to both stop-gain SNVs and fs-indels, as both trigger premature termination codons. Using the skin cutaneous melanoma (SKCM) TCGA cohort, we annotated all fs-indels (n=1,594) and stop-gain SNVs (n=9,883) for exonic position. Penultimate and last exon alterations were found to be significantly depleted in fs-indels compared to stop-gain events (OR=0.58 [0.46-0.71], P=1.5×10^−5^ and OR=0.65 [0.55-0.75], P=1.5×10^−7^ respectively) (**Fig. 4**). By contrast fs-indel mutations were more likely to occur in middle exon positions (OR=1.51 [1.33-1.68], P=1.2×10^−11^). First exon mutations were not enriched either way, possibly due to small absolute numbers (only n=69 fs-indels were first exon). This data suggests negative selective immune pressure acts against fs-indel mutations in exonic positions likely to escape NMD (e.g. penultimate and last), leading to cancer cells with middle exon fs-indels being more likely to survive immunoediting. As an additional control to rule out any potential bias in variant calling between fs-indels and stop-gain SNV groups, we repeated the above analysis for germline variants from the Exome Aggregation Consortium (ExAC) database (27). Due to self-tolerance no immunogenicity would expected against either stopgain or fs-indels, and in accordance with this no depletion in fs-indel mutations was detected in penultimate or last exon positions (all ORs were between 0.8 and 1.2) (**Fig. 4**). Finally, to assess evidence of natural anti-tumor immunogenicity of NMD-escape mutations in melanomas, we examined matched DNA and RNA sequencing data from 368 patients in the TCGA SKCM cohort. Patients with at least one NMD-escape mutation had significantly improved OS (HR=0.69 [0.50-0.96], P=0.03), as compared to those with zero NMD-escape mutations (**Fig. S2**). Additionally, using matched DNA and RNA sequencing data from a small cohort MSI carcinomas (which have high abundance of fs-indel events) identified by Cortes-Ciriano et al. (28) (n=96), a non-significant OS difference was observed among patients with high NMD-escape mutation load (defined as > cohort median value rather than =>1, due to the high level of indel events) (HR=0.67 [0.31-1.45], P=0.313) (**Fig. S2**).

**Figure 4.**
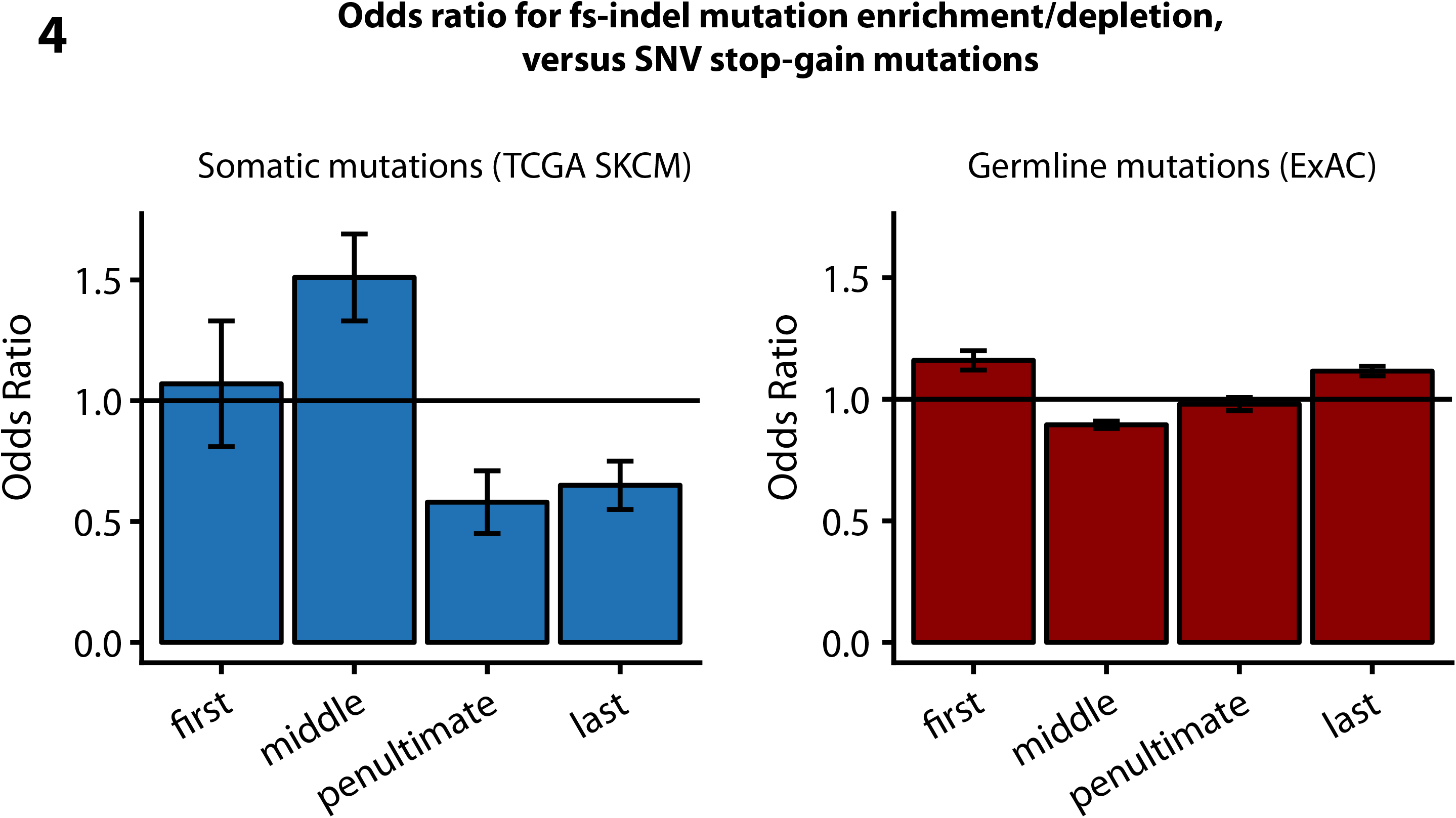
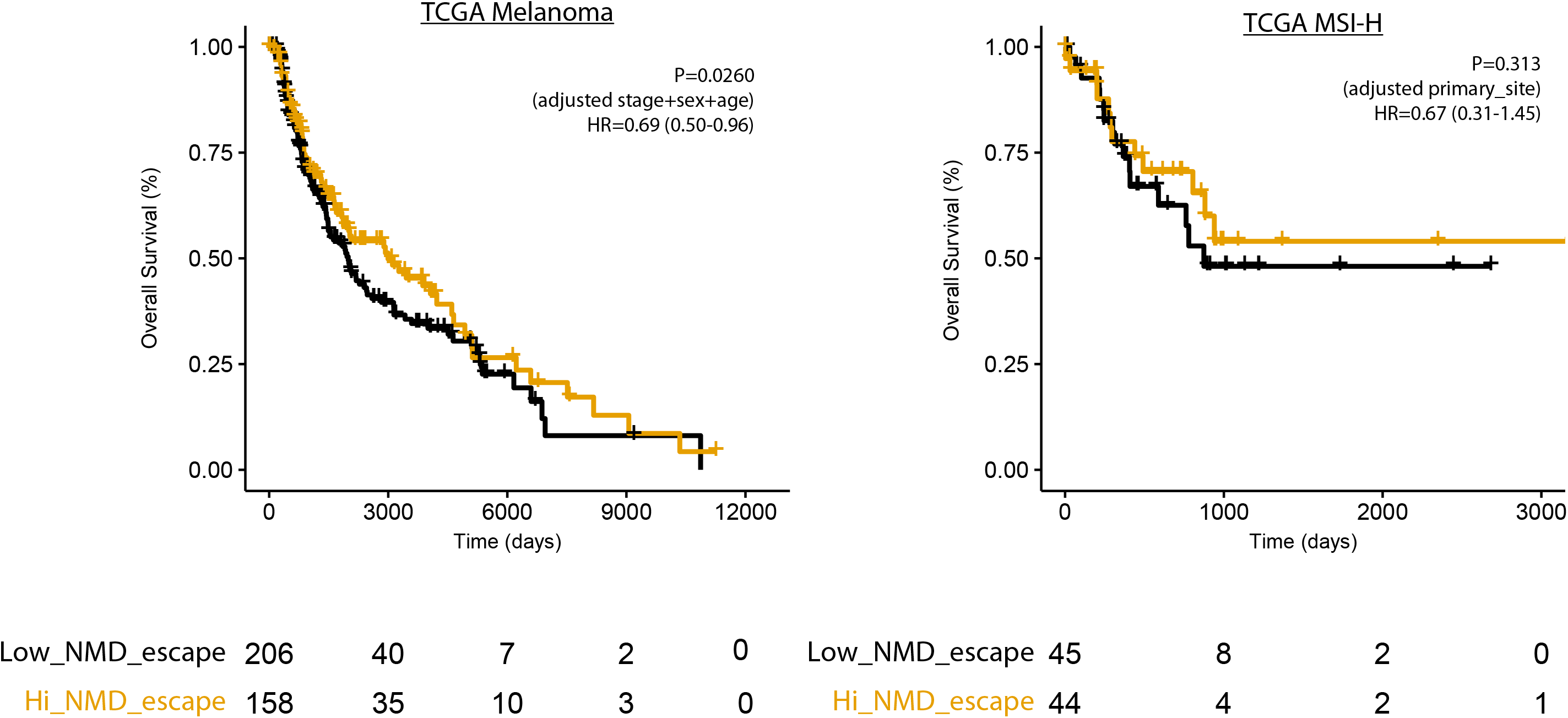
Shows selection analysis for fs-indels, as benchmarked against functionally equivalent SNV stop-gain mutations. The odds ratio for a fs-indel (compared to SNV stop-gains), to fall into each exon position group is shown. On the left is somatic mutational data from TCGA and on the right is germline mutation data (used as a negative control) from the ExAC database. Odds ratios and associated p-values were calculated using Fisher’s Exact Test. Coloring is used arbitrarily to distinguish groups.

## Discussion

In this study, we analysed expressed fs-indels, in the context of NMD and anti-tumor immunogenicity. We show that expressed fs-indels are highly enriched in genomic positions predicted to escape NMD, and have higher protein-level expression (relative to non-expressed fs-indels). Expressed fs-indels (a.k.a. NMD-escape mutations) also significantly associated with clinical benefit from immunotherapy. The small and heterogeneous individual cohorts (^~^20-30 cases/cohort) utilized in this study should be acknowledged however.

The primary controlling mechanism of NMD in mammalian cells is proposed to be the exon junction complex (EJC) model, whereby transcripts bearing a premature termination codon (PTC) >50bp upstream of the last exon junction escape degradation. Indeed, we observe strong consistency with this model in comparing expressed versus non-expressed fs-indels, with the former being highly enriched in penultimate and last exon position mutations (both OR>1.8, P<5×10^−12^). In addition, protein abundance was found to be significantly higher in the expressed fs-indel group (P=0.018). The unexplained determinants of NMD should also be recognised however, with recent studies (14) estimating that over a quarter of NMD variance remains unexplained across all genes. Similarly, these exceptions are visible in our data, with an appreciable number of mutations in middle exon position detected as expressed. As well as novel instances of NMD-escape, it is also possible a subset of these middle exon mutations are in fact undergoing partial (or full) NMD degradation, but remain at sufficiently high transcript abundance to be detected. Translational plasticity has also been described as additional feature impacting NMD efficiency, with a diverse range of mechanisms such as stop codon read through, alternative translation initiation and alternative splicing, being reported as driving NMD escape in the germline setting (29). Furthermore, the highly dysregulated nature of cancer cell transcriptomes may further explain the partial “leakiness” in NMD patterns we observe in this study. These exceptions, and the currently incomplete understating of NMD, highlights the importance of establishing fs-indel expression using RNA sequencing data and the need for further mechanistic research in this area.

NMD-escape mutation count was found to significantly associate with clinical benefit from immunotherapy, across both CPI and ACT modalities, and with a stronger association than either nsSNVs or fs-indels. CPI clinical benefit rates for patients with ≥ one NMD-escape mutation were elevated (range across the cohorts analysed = 0.56-0.75) compared to patients with zero such events (range 0.12-0.35). Furthermore, NMD-escape mutation count was shown to remain significantly associated with clinical benefit to CPI in the low-TMB setting (P=0.013), whereas nsSNV count was not (P=0.19). This raises the prespect of “rescuing” patients who may fall below the overall 10 mutations per megabase TMB threshold, but have a higher chance of CPI response based on harboring one or more NMD-escape mutation events. Several potential sources of antigenic peptide material for human leucocyte antigen presentation are proposed, ranging from classical degradation of previously functional proteins, to alternative sources including the pioneer round of mRNA translation and defective ribosomal products (30) (31). The enrichment of NMD-escape features, in the expressed fs-indels observed in this study, would favour a classical route of translational as at least one source of peptide material in these cohorts. However, the appreciable number expressed fs-indels deriving from middle exon positions, suggests additional possible sources may be present, such as those from the pioneer round of translation.

Experimental evidence, analyzed from anti-tumor vaccine and CPI studies, demonstrates T cell reactivity against expressed frameshifted neoepitopes directly in human patients (n=15). T cell reactive fs-indel neoantigens were also enriched for longer neoORF length (median=27 amino acids), versus experimentally screened, but T cell non-reactive fs-indels (median=5 amino acids) (P=0.0032). These elongated neoORF mutations create the additional benefit of increased redundancy in HLA allele binding, based on the intuiative result that a greater number of peptides will be capable of binding to a broader spectrum of patient HLA-alleles. Selection analysis demonstrated a depletion of fs-indels in penultimate and last exon positions, as compared to functionally equivalent stop-gain SNVs, suggesting potential negative immune selection against NMD-escape events during tumor evolution. As a negative control, we demonstrate this same association is not present in germline mutations from the ExAC database. Checkpoint control of immune response is a likely compensatory mechanism used by tumor cells to manage remaining NMD-escape events, a notion in keeping with the elevated CPI clinical benefit rates we see in patients with even a small number of NMD-escape alterations. NMD-escape count was found to be prognostic in untreated TCGA melanoma cases, consistent with the hypothesis of NMD-escape mutations driving native immunity. We did not observe an association between NMD-escape mutation count and CPI clinical benefit in a metastatic urothelial cancer cohort. We note that in this cohort neither TMB, predicted neoantigen nor expressed neoantigen load were associated with clinical benefit in the original report (20), perhaps due to limited statistical power (n=23 cases). In terms of study limitations, we acknowledge that escape of NMD has not been functionally demonstrated in this work, and instead the enrichment of penultimate/last exon position mutations, together with higher protein expression levels, is suggestive of NMD-escape however not direct proof. This limitation is in keeping with the translational biomarker scope of this study, however further functional investigation of NMD-escape mechanisms will be of significant interest. Furthermore, we acknowledge that neoantigen presentation is an inefficient process, and that NMD-escape mutations identified by DNA/RNA sequencing are unlikely to directly causative in all tumors. Here we present evidence to support that NMD-mutations as a rare mutation type, with enhanced potential to elicit an anti-tumor immune response. A further limitation are the bioinformatics challenges in accurate indel calling, meaning a reduced sensitivity to detect all fs-indels. However, here we demonstrate that using both DNA and RNA sequencing assays improves calling accuracy and leads to a high confidence call set, due to alteration detection at both DNA and RNA levels (see methods). In summary, here we highlight NMD-escape mutations as a highly immunogenic mutational subset, rare in nature but found to significantly associate with clinical benefit to immunotherapy. These mutations may represent attractive targets for personalized immunotherapy design, as well as contributing to the refinement of genomic biomarkers to predict CPI response.

## Materials and methods

### Study cohorts

Matched DNA/RNA sequencing analysis was conducted in the following cohorts all treated with immunotherapy:

- Van Allen et al. (8), an advanced melanoma checkpoint inhibitor (CPI) (anti-CTLA-4) treated cohort. Cases with both RNA sequencing and whole exome (DNA) sequencing data were utilised (n=33).
- Snyder et al. (7), an advanced melanoma CPI (anti-CTLA-4) treated cohort. Cases with both RNA sequencing and whole exome (DNA) sequencing data were utilised (n=21).
- Hugo et al. (4), an advanced melanoma CPI (anti-PD-1) treated cohort. Cases with both RNA sequencing and whole exome (DNA) sequencing data were utilised (n=24).
- Riaz et al. (32), an advanced melanoma CPI (anti-PD-1) treated cohort. Cases with both RNA sequencing and whole exome (DNA) sequencing data, from the ipilimumab-naive cohort, were utilised (n=24). In keeping with the original publication, we found the other patient-cohort in this study (cases pre-treated and progressive on ipilimumab therapy (Ipi-P)), to have no association between mutation load metrics (nsSNVs, fs-indels, NMD-escape mutations) and subsequent benefit from anti-PD1 therapy.
- Lauss et al. (10), an advanced melanoma adoptive cell therapy treated cohort. Cases with both RNA sequencing and whole exome (DNA) sequencing data were utilised (n=22).
- Snyder et al. (20), a metastatic urothelial cancer CPI (anti-PD-L1) treated cohort. Cases with both RNA sequencing and whole exome (DNA) sequencing data were utilised (n=23).

Matched DNA/RNA sequencing analysis was conducted in the following cohorts (not specifically treated with immunotherapy):

- Skin cutaneous melanoma (SKCM) tumors, obtained from the cancer genome atlas (TCGA) project. Cases with paired end RNA sequencing data and curated variant calls from TCGA GDAC Firehose (2016_01_28 release) were utilised (n=368).
- Microsatellite instable (MSI) tumors, across all histological subtypes from TCGA project. MSI cases IDs were identified based on classification from Cortes-Ciriano et al. (28). Cases with paired end RNA sequencing data and curated variant calls from TCGA GDAC Firehose (2016_01_28 release) were utilised (n=96).

Prediction of NMD-escape features (based on DNA exonic mutation position only, rather than matched DNA/RNA sequencing analysis) was conducted in the following immunotherapy treated cohorts:

- Ott et al. (21), an advanced melanoma personalized vaccine treated cohort (n=6 cases).
- Rahma et al. (22), a metastatic renal cell carcinoma personalized vaccine treated cohort (n=6 cases).
- Le et al. (23), an advanced mismatch repair-deficient cohort, across cancers across 12 different tumor types, treated with anti-PD-1 blockade (n=86 cases, functional neoantigen reactivity T cell work only conducted in n=1 case).

### Whole exome sequencing (DNA) variant calling

For Van Allen et al. (8), Snyder et al. (7) and Snyder et al. (20) cohorts, we obtained germline/tumor BAM files from the original authors and reverted these back to FASTQ format using Picard tools (version 1.107) SamToFastq. Raw paired-end reads in FastQ format were aligned to the full hg19 genomic assembly (including unknown contigs) obtained from GATK bundle (version 2.8), using bwa mem (bwa-0.7.7). We used Picard tools to clean, sort and to remove duplicate reads. GATK (version 2.8) was used for local indel realignment. We used Picard tools, GATK (version 2.8), and FastQC (version 0.10.1) to produce quality control metrics. SAMtools mpileup (version 0.1.19) was used to locate non-reference positions in tumor and germline samples. Bases with a Phred score of less than 20 or reads with a mapping quality less than 20 were omitted. VarScan2 somatic (version 2.3.6) used output from SAMtools mpileup to identify somatic variants between tumour and matched germline samples. Default parameters were used with the exception of minimum coverage for the germline sample, which was set to 10, and minimum variant frequency was changed to 0·01. VarScan2 processSomatic was used to extract the somatic variants. Single nucleotide variant (SNV) calls were filtered for false positives with the associated fpfilter.pl script in Varscan2, initially with default settings then repeated with min-var-frac=0·02, having first run the data through bam-readcount (version 0.5.1). MuTect (version 1.1.4) was also used to detect SNVs, and results were filtered according to the filter parameter PASS. In final QC filtering, an SNV was considered a true positive if the variant allele frequency (VAF) was greater than 2% and the mutation was called by both VarScan2, with a somatic p-value <=0.01, and MuTect. Alternatively, a frequency of 5% was required if only called in VarScan2, again with a somatic p-value <=0.01. For small scale insertion/deletions (INDELs), only calls classed as high confidence by VarScan2 processSomatic were kept for further analysis, with somatic_p_value scores less than 5 × 10^−4^. Variant annotation was performed using Annovar (version 2016Feb01). For the Hugo et al. (4) cohort, we obtained final post-quality control mutation annotation files generated as previously described (4). Briefly, SNVs were detected using MuTect, VarScan2 and the GATK Unified Genotyper, while INDELs were detected using VarScan2, IndelLocator and GATK-UGF. Mutations that were called by at least two of the three SNV/INDEL callers were retained as high confidence calls. For the Lauss et al. (10) cohort, SNVs and INDELs were called as described previously (10). Briefly, SNVs were detected using the intersection of MuTect and VarScan2 variants, while INDELs were detected using VarScan2 only. For VarScan2, high confidence calls at a VAF greater than 10% were retained.

### Whole transcriptome sequencing (RNA) variant calling

RNAseq data was obtained in BAM format for all studies, and reverted back to FASTQ format using bam2fastq (v1.1.0). Insertion/deletion mutations were called from raw paired end FASTQ files, using mapsplice (v2.2.0), with sequence reads aligned to hg19 genomic assembly (using bowtie pre-built index). Minimum QC thresholds were set to retain variants with => 5 alternative reads, and variant allele frequency => 0.05. Insertions and deletions called in both RNA and DNA sequencing assays were intersected, and designated as expressed indels, with a +/- 10bp padding interval included to allow for minor alignment mismatches. SNVs in RNA sequencing data were called directly from the hg19 realigned BAM files, using Rsamtools to extract read counts per allele for each genomic position where a SNV had already called in DNA sequencing analysis. Similarly, minimum QC thresholds of => 5 alternative reads, and variant allele frequency => 0.05, were utilised and variants passing these thresholds were designated as expressed SNVs.

### Consensus indel variant calling accuracy

As an additional methodological check, indels were re-called from datasets (7) and (8), using two additional DNA variant callers, Mutect2 and Scalpel, in addition to Varscan2. The aim was to assess if the joint DNA and RNA calling approach used in this study lead to higher consensus between variant callers, and hence a reduction in the risk of caller specific artefacts. Using a DNA calling only approach, we observed the consensus between variant callers ranging from 67% [called in all three tools Varscan2/Mutect2/Scalpel] to 82% [called in Varscan2 and one of Mutect2 or Scalepl]. These same values for indels called in both DNA and RNA sequencing data increased to 81% and 100% respectively. Thus, we find 100% of the reported NMD-escape indel mutations (detected in both DNA and RNA) were called in two or more different DNA indel calling algorithms.

### Isoform annotation

For analysis in figure 1B and 1C, variants were annotated on a tumor specific isoform basis. Specifically, level 3 isoform obtained was obtained from the broad firehose repository (https://gdac.broadinstitute.org/). For each mutated gene in each tumor, the corresponding isoform expression values were extracted (for the gene in question), and the isoform with highest adbudance was selected for annotation purposes. Isoform annotation was conducted using Annovar, with frameshift indel mutations grouped into five categories, based on the position of the premature termination codon following the frameshift: i) first exon, ii) middle exon, iii) penultimate exon more than 50bp of the last exon junction complex, iv) penultimate exon less than or equal to 50bp of the last exon junction complex, v) last exon.

### Protein expression analysis

We retrieved Level 4 (L4) normalized protein expression data for 223 proteins, across n=453 TCGA melanoma/MSI tumors (which overlapped with the TCGA cohorts also analysed via DNA/RNA sequencing) from the cancer proteome atlas (http://tcpaportal.org/tcpa/index.html). We filtered the data to sample/protein combinations which also contained an fs-indel mutation (n=136), as called by DNA sequencing. The dataset was then split into two groups, based on the fs-indel being expressed or not (as measured by RNAseq, using the method detailed above). The two groups were compared using a two-sided Mann Whitney test.

### Outcome analysis

Across all immunotherapy treated cohorts, measures of patient clinical benefit/no-clinical benefit were kept as consistent with original author’s criteria/definitions. For TCGA outcome analysis, overall survival (OS) data was utilized, based on clinical annotation data obtained from TCGA GDAC Firehose repository.

### Selection analysis

To test for evidence of selection, fs-indel mutations were compared to stop-gain SNV mutations, in the SKCM TCGA cohort (n=368 cases). Stop-gain SNV mutations were utilised a benchmark comparator, due to their likely equivalent functional impact (*i.e*. loss of function), equivalent treatment by the NMD pathway (*i.e*. last exon stop-gain SNVs will still escape NMD and cause truncated protein accumulation) but lack of immunogenic potential (*i.e*. no mutated peptides are generated). Across all SKCM cases n=1,594 fs-indels and n=9,833 stop-gain SNVs were considered. All alterations in each group were annotated for exon position (*i.e*. first, middle, penultimate or last exon, as defined above). The odds of having an fs-indel in first, middle, penultimate or last exon positions was then benchmarked against the equivalent odds for a stop-gain SNV.

### Statistical methods

Odds ratios were calculated using Fisher’s Exact Test for Count Data, with each exon position group compared to all others. Kruskal-Wallis test was used to test for a difference in distribution between three or more independent groups. Two-sided Mann Whitney U test was used to assess for a difference in distributions between two population groups. Meta-analysis of results across cohorts was conducted using the Fisher method of combining *P* values from independent tests. Logistic regression was used to assess multiple variables jointly for independent association with binary outcomes. Overall survival analysis was conducted in the SKCM TCGA cohort using a Cox proportional hazards model, with stage, sex and age included as covariates. Overall survival analysis was conducted in the MSI TCGA cohort using a Cox proportional hazards model, with primary disease site included as a covariate. Statistical analysis were carried out using R3.4.4 (http://www.r-project.org/). We considered a P value of 0.05 (two sided) as being statistically significant.

## Author’s contributions

Study design: K.L., S.T., C.S

Data analysis: K.L., E.L., S.T.

Data interpretation: all authors

Manuscript writing: K.L., M.D.H, S.T, C.S

## Table Legends

**Table S1**

Screened neoORF mutations from human studies.

